# Systematic comparative analysis of Mendelian randomization methods for inferring causal genes of complex phenotypes and the application to psychiatric diseases

**DOI:** 10.1101/2020.11.09.374298

**Authors:** Lin Jiang, Guorong Yi, Xiangyi Li, Chao Xue, Mulin Jun Li, Hailiang Huang, Miaoxin Li

**Affiliations:** Research Center of Medical Sciences, Guangdong Provincial People’s Hospital, Guangdong Academy of Medical Sciences, Guangzhou, China; Zhongshan School of Medicine, Sun Yat-sen University, Guangzhou, 510080, China; Key Laboratory of Tropical Disease Control (Sun Yat-sen University), Ministry of Education, Guangzhou, 510080, China; Center for Precision Medicine, Sun Yat-sen University, Guangzhou, 510080, China; Medical University Cancer Institute and Hospital, Tianjin Medical University, Tianjin 300070, China; The Province and Ministry Co-sponsored Collaborative Innovation Center for Medical Epigenetics, Tianjin; Massachusetts General Hospital, 185 Cambridge St, Boston MA 02114; State Key Laboratory of Brain and Cognitive Sciences, the University of Hong Kong, Pokfulam, Hong Kong

**Keywords:** Mendelian randomization, causal gene, expression quantitative trait locus, genome-wide association study

## Abstract

Isolating causal genes from enormous genome-wide association signals of complex phenotypes remains an open and challenging question. SMR (Summary-based Mendelian Randomization) is a widely used Mendelian randomization (MR) method for inferring causal genes by using a single expression quantitative trait locus (eQTL). In the present study, we explored more powerful MR methods based on multiple eQTLs. Among six representative multiple instrumental variable (IVs) based MR methods, original used in the epidemiological field, not all MR methods worked for the causal gene estimation. But we found the maximum-likelihood based MR method and weighted median-based MR method were preferable to the other four MR methods in terms of valid type 1 errors, acceptable statistical powers and robustness to linkage disequilibrium (LD) in eQTLs. Both of the MR methods were also much more powerful than the SMR. We recalibrated key parameters of the two MR methods in practices and developed a multiple IVs based MR analysis framework for causal gene estimation, named MACG and available at http://pmglab.top/kggsee. In the applications, MACG not only rediscovered many known causal genes of the schizophrenia and bipolar disorder, but also reported plenty of promising candidate causal genes. In conclusion, this study provided a powerful tool and encouraging exemplars of mining potential causal genes from huge amounts of GWAS signals with eQTLs.

## Introduction

While large scale genome-wide association studies (GWAS) have found huge amounts of single nucleotide polymorphism (SNP) loci associated with complex phenotypes, it still remains a challenge to identify causal genes from numerous phenotype-associated variants^1^. Genotype-phenotype association analysis is a crude investigation in the complex biological process transmitting the DNA information to transcribed RNA, then to translated proteins and finally to phenotypes ^2 3^. Notably, most of trait-associated variant are located in non-protein coding regions ^4^ and do not affect composition of proteins. These non-coding variants often imply regulation of gene expression, felicitously explaining the mechanism from genotypes to phenotypes. Therefore, many transcription resources have been released to motivate researchers to decipher genetic mechanism of complex phenotypes, including multiple large collaborative genes expression resources, such as GTEx^4^, CommonMind Consortium^5^, and GEUVADIS^6^.

To make the best of these valuable and increasing gene transcription resources, advanced methods are urgently needed to infer causal paths according to gene expression regulation. Many methods are being developed to link gene expression to phenotypes. Others’ and our recent studies also showed that phenotype-associated genes tend to have selected expression in phenotype related tissues or cell-types; and gene’s selective expression was convincing evidence for prioritizing phenotype-associated genes and related tissues^7,8^. Some methods have used expression quantitative trait loci (eQTLs) analysis as an auxiliary instrument to interpret GWAS signals because GWAS hits are found to be enriched with eQTL^9^ and being eQTLs are prerequisite for the loci to regulate a phenotype by gene expression^10 9^. Directly based on the top cis-eQTLs, Zhu et al proposed a method, named SMR (Summary-based Mendelian Randomization), to infer the causal pathway from genotype to transcription and then to phenotype^11^. As a single eQTL usually has limited power to detect susceptibility genes, Gusev et al built a framework known as PrediXcan, which used multiple eQTLs to predict gene expression and then examined the association between the imputed expression and a phenotype ^12^. This framework was further extended to GWAS summary statistics, named S-PrediXcan^13^, which is flexible for large-scale GWAS samples. However, except for SMR, all of these methods with multiple SNPs are designed for detecting association rather than the causation between gene expression and complex phenotypes.

Mendelian randomization (MR) is a widely used strategy to infer causal relationship between an exposure and an outcome^14^. Basically, it assumes that a genotype should be correlated to the outcome when genetic variants (as instrumental variables, IVs) alter the level of a modifiable exposure which causally changes the outcome. The causal effect of exposure on outcome is equal to the correlation between genotype and outcome divided by the influence of genotype on exposure^15^. Recent MR methods used two samples to infer the causality between phenotypes with multiple IVs^16^ in epidemiological fields. As the path from genetic variants to exposure and to outcome may also be distorted by confounding factors (e.g. variants with pleiotropy), several MR methods were proposed to decrease influence of the confounding factors (e.g., MR-Egger^17^ and contamination mixture MR^18^). To our knowledge, in genomics field, no multiple IVs based MR methods has been applied to infer causal gene expression of complex phenotypes. So, it is worth more attention to pay on the unique practical concerns in transferring the multiple IV based MR methods to adapt this new field. First, the sample sizes of exposures and outcomes are usually very unbalanced. While GWAS summary statistics are usually generated from large-scale samples (say, n>10,000), the eQTLs are calculated from much smaller samples (e.g. several hundred subjects or even fewer in GTEx^4^). Second, the heritability of the cis-variants for gene expression (∼0.2) and complex phenotypes (<0.01) is also very unbalanced^19^. Third, different with the conventional applications of MR methods, only a small number of genetic variants are available in eQTLs results, say, ∼10-100 cis-eQTLs near a gene. By contrast, in conventional applications, variants for the causality inference are numerous, even from the entire genome.

We hypothesized that some of existing multiple IVs based MR methods can be recalibrated to infer causal genes with the unbalanced samples. The present study aims to develop a more powerful framework to infer causal genes of phenotypes with GWAS summary statistics and eQTLs. Within systematic simulations and real data validations, we prioritized suitable MR methods from six representative MR methods and recalibrate key parameters to address the above new technical problems in mining causal genes from GWAS association signals. The methods were inverse-variance weighted (IVW) MR^20^, maximum-likelihood (ML) based MR^16^, MR-Egger method^17^, median-based MR^21^, mode-based MR^22^ and contamination mixture MR^18^. Based on the investigation, a framework with abundant eQTL resource and the best practice pipeline were then developed to detect potential causal genes of complex phenotypes.

## Results

### Overview of the Mendelian randomization framework for inferring causal genes of complex phenotypes

Mendelian randomization is conventionally developed to estimate causal relationship among epidemiological phenotypes. Here we proposed a framework of Mendelian randomization analysis for causal gene inference, named MACG, with GWAS summary statistics and eQTLs (See workflow in Figure 1B). Figure 1A shows the overview of the assumption. Assume a number of variants (Z) regulate or associate with gene expression(X, exposure) in a relevant tissue of a phenotype (Y, outcome), as the expression subsequently contributes to the development of the phenotype. In addition, some other variants may directly regulate the expression and the phenotype simultaneously, usually called pleiotropic variants. Meanwhile, a number of confounding factors (U), which increase the correlation (but not causality) between expression and phenotypes, also regulate the expression and phenotype.

**Figure 1.**
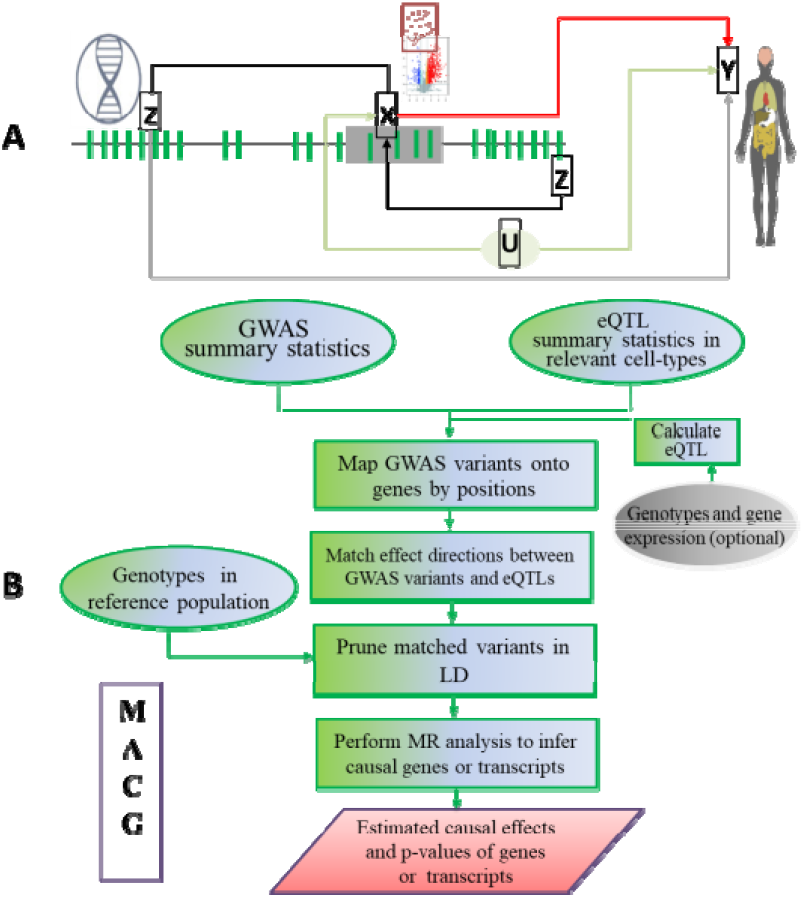
The diagram and workflow of the framework for inferring causal effects of gene expression on a phenotype. A) the diagram. B) the workflow of the framework. Z denotes IVs of MR. X denotes expression of transcripts as an exposure of MR and Y denotes a complex phenotype as the outcome of MR. The gene expression directly regulates phenotypes. The complex phenotype and expression are also regulated by confounding factors (U). In addition, some instrument variables also directly regulate both the expression and phenotype (known as pleiotropy effect). The proposed approach aims to infer causal effect from X to Y.

MACG adopted two multiple IVs based MR methods for causality test and casual effect estimation of a gene’s expression to a phenotype, median-based MR and ML-based MR. These are two selected methods out of six MR methods with recalibrated key parameters according to type 1 errors, statistical power, estimation accuracy, sensitivity to pleiotropy and correlation in IVs. The six MR methods had different representative designs to address issues of horizontal pleiotropy or correlation in IVs (see detailed descriptions about the MR methods in Methods and Materials section). Figure 1B shows the workflow and major datatypes. MACG needs two major inputs, GWAS and eQTL summary statistics respectively. The GWAS summary statistics refer to the logarithm of odds ratio or regression coefficients and the corresponding standard errors (SEs) from a large-scale GWAS study, indicating the association between IVs and a phenotype. The eQTL summary statistics are similar to that of the GWAS, indicating association between IVs and expression of genes or transcripts in a tissue or cell type. To avoid weak IV bias^23^, eQTLs with p≥1E-4 are excluded. MACG has integrated the pre-calculated cis-eQTLs in 55 tissues or cell-types with gene-level and transcript-level expression from GTEx^24^ (version 8). MACG can also use local expression to generate eQTLs on MACG. MACG maps the input variants onto genes with 1 MB window expansion away from the original gene boundaries on both sides. To ensure the allele coding is concordant between the GWAS variants and eQTL, MACG also checks the frequencies the coded alleles. The sign of coefficients will be flipped when the frequency difference suggests reversed coding. Using genotypes in reference populations, MACG pruned variants in high LD (*r*>0.5, see reasons for this cutoff in the following section). Finally, the coefficients and SEs of the IVs are analyzed by the MR methods to estimate and test causal effects of a gene or transcript to a phenotype. MACG has been implemented into our software platform KGGSEE, available at http://pmglab.top/kggsee.

### Type 1 error of six MR methods for identifying causal genes

We first systematically evaluated the type 1 error rate of six widely-used MR methods for inferring causal genes by multiple correlated eQTLs. We explored three scenarios about IVs for both continuous and binary phenotypes (heritability): an ideal scenario for MR model, a scenario with confounding factors (and), and a scenario with pleiotropy (and). Three levels of total eQTL heritability (=0.05, 0.15 and 0.25) were also investigated for each scenario. For a continuous phenotype, the median-based MR method and contamination mixture MR method showed inflated type 1 errors under the null hypothesis that a gene has no causal effect on the phenotype. The mode-based MR showed slight inflation when the heritability of eQTLs was as high as 0.25 in the simulated datasets. The remaining three MR methods, considering LD of IVs, showed reasonable type 1 errors (Figure 2). It should be noted that in the pleiotropy scenario half of IVs were invalid due to direct pleiotropic effect on gene expression and phenotypes (Figure 2c, 2f, 2i). MR-Egger method was designed to correct for pleiotropy ^25^ which could explain its validity of type 1 error in this scenario. Although the IVW MR method and the ML-based MR method did not specifically account for pleiotropy, both methods under random-effects models also worked well in the simulations. For binary phenotypes, all of the type 1 error patterns observed in the continuous traits were replicated regardless of whether the odds ratios were generated by logistic regression model (Figure S1) or by conventional contingency table analysis (Figure S2).

**Figure 2.**
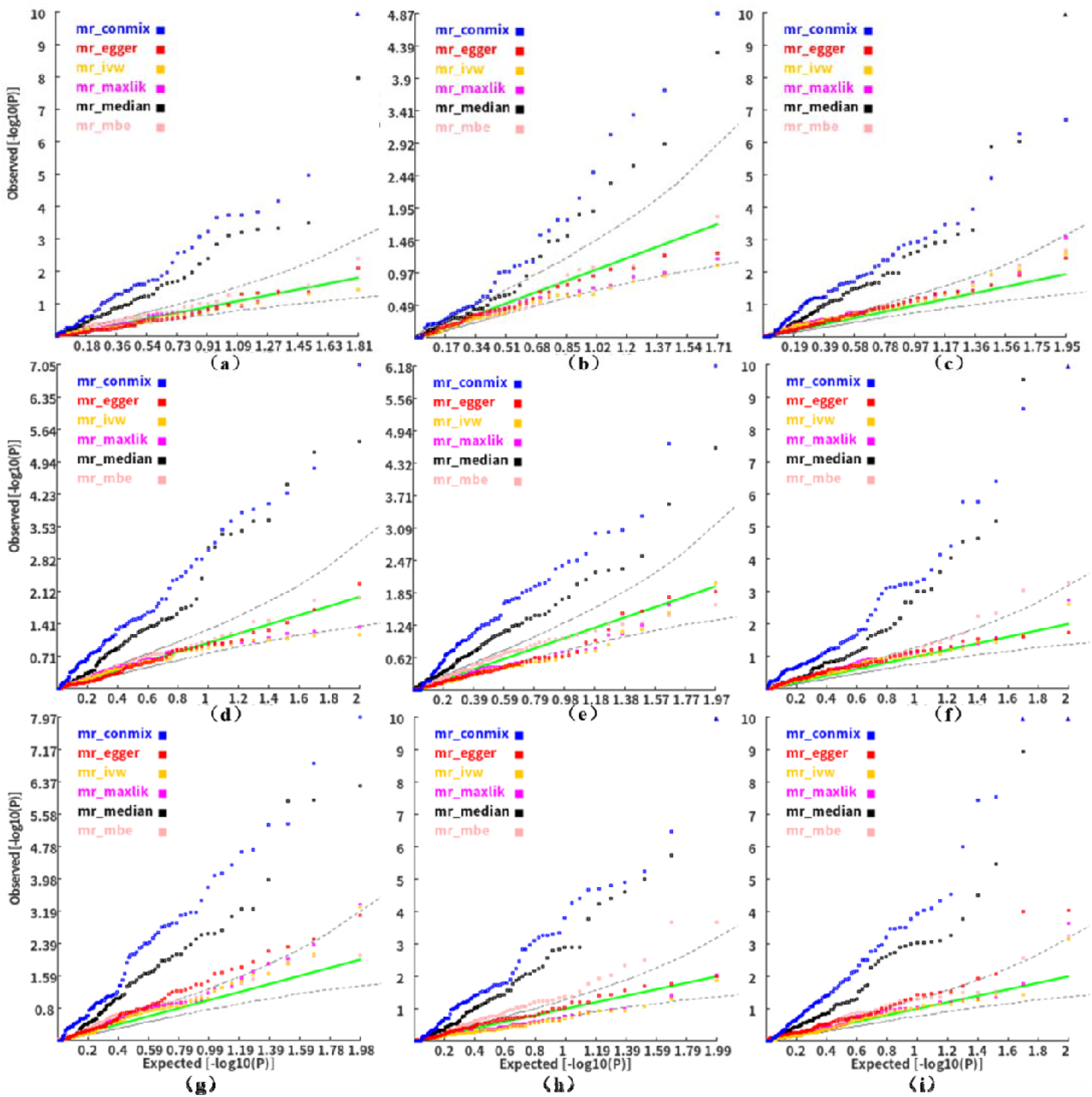
Type 1 errors of six MR methods for inferring causal genes of a continuous phenotype. a), d) and g) show the type 1 error of MR methods under an ideal scenario with total eQTL heritability 0.05, 0.15 and 0.25 respectively. b), e) and h) show the type 1 error of MR methods under a scenario with confounding factors with total eQTL heritability 0.05, 0.15 and 0.25 respectively. c), f) and i) show the type 1 error of MR methods under a scenario with pleiotropy with total eQTL heritability 0.05, 0.15 and 0.25 respectively.

As the inflated type 1 error in the three above MR methods might be caused by correlation in the IVs, we pruned the SNPs according to various LD thresholds to investigate the type 1 errors of the six MR methods again. For a continuous phenotype, when the LD threshold was high (say, *r*=0.9), all the five MR methods except for mode-based estimation (MBE)MR showed inflated type 1 error rate (Figure S3). Unlike the preceding analysis, the IVW MR method and the ML-based MR method did not consider correlation of IVs here. As the LD-pruning threshold decreased to a moderate level, say *r*=0.5, the type 1 error of two MR methods (i.e. ML-based MR and median-based MR) became valid approximately. MR-Egger showed the highest inflation in type 1 error while the MBE MR method became deflated (Figure S3). For a binary phenotype, the MR methods showed similar sensitivity to LD as they did for the continuous phenotype. When the LD-pruning threshold *r* was 0.5, the ML-based MR method and median-based MR method also showed valid type 1 error approximately (Figure S4). Other patterns of type 1 errors of the six MR methods based on the LD-pruned IVs for a binary phenotype were similar to that of a continuous phenotype.

We also investigated how pleiotropic effects influenced the type 1 errors of the six MR methods. It turned out the ML-based MR and median-based MR methods were also insensitive to pleiotropy with moderately LD-pruned IVs (Figure S17). For example, when the heritability values of pleiotropic variants to a continuous phenotype and gene expression were 0.03 and 0.15, the two methods still produced approximately uniformly distributed p-values. There were no obvious inflation even when the heritability values of pleiotropic variants to the phenotype were as high as 0.05. MR-Egger showed the highest inflation in type 1 error regardless of degree of pleiotropy. The contamination mixture MR and IVW MR showed slight inflation when there was pleiotropy. The MBE MR was slightly deflated no matter what degree the pleiotropy was. These pleiotropic patterns of binary phenotypes were similar to that of the continuous phenotypes.

Therefore, the systematic simulations suggested three methods with correlation of IVs (IVW MR, ML-based MR, and MR-Egger) were valid to test gene’s causal expression to a phenotype by cis-eQTLs in terms of type 1 error. Meanwhile, ML-based MR and median-based MR were also valid approximately for the test when IVs were only moderately correlated (*r*<0.5) regardless of a continuous or binary phenotype.

### Statistical power comparison of MR approaches to infer causal effect of genes

We then investigated statistical power of the above MR methods with valid type 1 errors under their corresponding scenarios. Using correlated IVs, we compared p-values of three methods for inferring causal gene expression to a phenotype, namely, IVW MR, ML-based MR, and MR-Egger. We found MR-Egger almost always produced less significant p-values than the other two MR methods in all scenarios (See p-value distributions in Figure S5 in continuous phenotypes and Figure S6-7 in binary phenotypes). When the causal effect was as low as 0.05, both IVW MR method and ML-based MR method had low power with the ideal IVs because it was hard to obtain significant true eQTLs for the calculation in the small expression sample. As expected, when the causal effect of expression on phenotype increased, the power of the MR methods increased accordingly. In our simulations, when the causal effect was 0.15, the power of IVW MR method and ML-based MR method became 80% for continues phenotype with 15000 subjects and expression sample of 500 subjects. The patterns of power in binary phenotypes were similar to that of the continuous trait regardless of whether the odds ratio and SE were generated by a logistic regression model or by a contingency table analysis (Figure S6 and S7 respectively).

With moderately LD-pruned IVs (*r*<0.5), we also investigated the power of the two MR methods with valid type 1 errors (i.e., ML-based MR and median-based MR) for continuous phenotypes and binary phenotypes respectively. For continuous phenotypes, the two methods had similar power when the size of causal effect was moderate, say 0.2∼0.6 (Figure 3). However, when the causal effect size was 0.1, the ML-based MR method produced smaller p-values than the median-based MR method in most simulated datasets, suggesting higher power of the former. Interestingly, median-based MR method became more powerful than ML-based MR method when the causal effect size was 0.7. The binary phenotypes seemed to lead to simpler patterns of power than the continuous phenotypes. In addition, median-based MR method tended to produce smaller p-values than the ML-based MR method regardless of the causal effect size (Figure 3). When the causal effect was 0.3, the p-values’ difference of the two methods seemed even larger than that of 0.5 and 0.7.

**Figure 3:**
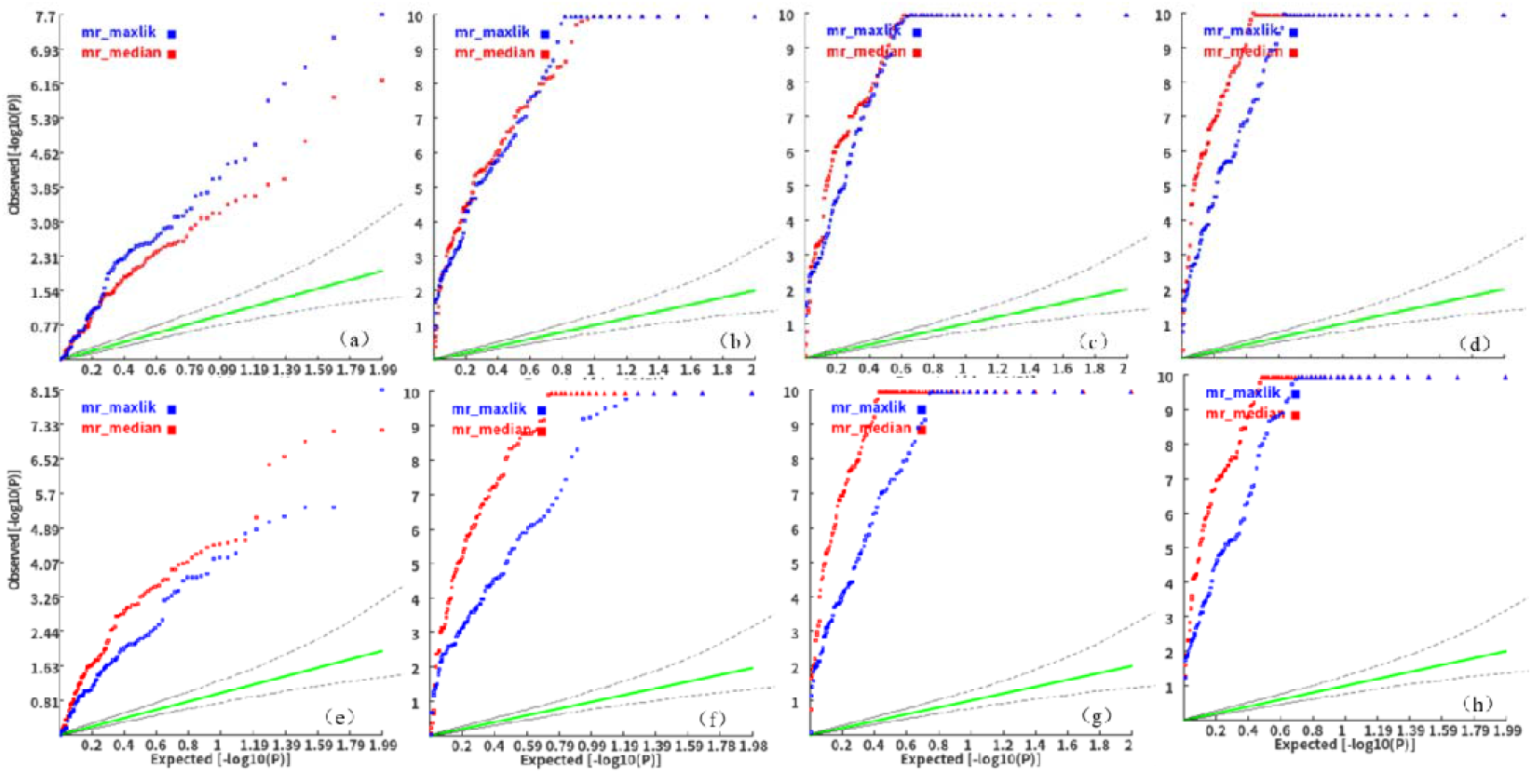
P-value distribution of four MR methods for inferring causal genes with LD-pruned IVs. The first, second, third and fourth columns correspond to the cause effects 0.1, 0.3, 0.5, and 0.7 respectively. The first and second rows correspond to continuous phenotype and a binary phenotype. The total heritability by eQTLs as IVs was 0.15.

### Compare the power of multiple IVs based MR method with SMR

To testify whether a multiple-IV based MR method is more powerful than that based on a single-IV, we finally compared the power of the ML-based MR method and Median-based MR method with SMR^11^. SMR uses a leading eQTL IV to test causal gene expression of a phenotype, which has been widely used in genetic field. It turned out that both of the multiple IV based MR methods were much more powerful than the single IV based MR method generally (Table 1). For continuous phenotypes, the ML-based MR method and Median-based MR method with LD-pruned IVs (*r*<0.5) achieved around 30% more power than the SMR when the causal effects were over 0.3 and there was no horizontal pleiotropy. When there was horizontal pleiotropy, SMR had some improved power to detect a causal gene. However, it was still ∼10-20% less powerful than the two multiple IV based MR methods. For binary phenotypes, the two multiple-IV based MR methods were also much more powerful than SMR. For example, when the causal effect was 0.3, the Median-based MR method with LD-pruned IVs (*r*<0.5) had 55% more power than the SMR by IVs without horizontal pleiotropy. When there was horizontal pleiotropy, the SMR had also enhanced power but was still less powerful than both multiple-IV based MR methods (See details in Table 1).

**Table 1:** Power of ML-based MR method, Median-based MR method and SMR for causal gene estimation

| Phenotype | H.P.V | Causal effects | ML-based MR | Median-based MR | SMR |
| --- | --- | --- | --- | --- | --- |
| continuous | 0 | 0.1 | 0.19 | 0.09 | 0.00 |
| continuous | 0 | 0.3 | 0.66 | 0.72 | 0.14 |
| continuous | 0 | 0.5 | 0.78 | 0.82 | 0.39 |
| continuous | 0 | 0.7 | 0.76 | 0.87 | 0.32 |
| continuous | 0.01 | 0.1 | 0.19 | 0.12 | 0.06 |
| continuous | 0.01 | 0.3 | 0.68 | 0.74 | 0.33 |
| continuous | 0.01 | 0.5 | 0.76 | 0.91 | 0.56 |
| continuous | 0.01 | 0.7 | 0.91 | 0.97 | 0.73 |
| binary | 0 | 0.1 | 0.37 | 0.31 | 0.04 |
| binary | 0 | 0.3 | 0.65 | 0.83 | 0.28 |
| binary | 0 | 0.5 | 0.68 | 0.80 | 0.39 |
| binary | 0 | 0.7 | 0.79 | 0.84 | 0.51 |
| binary | 0.01 | 0.1 | 0.20 | 0.33 | 0.23 |
| binary | 0.01 | 0.3 | 0.72 | 0.85 | 0.54 |
| binary | 0.01 | 0.5 | 0.80 | 0.94 | 0.68 |
| binary | 0.01 | 0.7 | 0.89 | 0.96 | 0.67 |

Each power was calculated from 100 simulated datasets. The details of simulating genotypes and phenotypes can be seen in the results section. The p-value threshold for a significant causality test was 0.001. H.P.V.: Heritability of pleiotropic variants

### Estimation accuracy for genes’ causal effect

After above performance evaluation for hypothesis test, we then asked whether the two valid and powerful MR methods with full set of IVs can accurately estimate the causal effect by more simulations. For continuous phenotypes, it turned out that the absolute value of estimated effects was generally smaller than that of the true effects (See Figure 4a, S8a and S8b). Note this was a non-conventional application of the MR methods where the size of expression samples was very small, equal to ∼3% of the GWAS samples, in our simulation studies. The sample size ratio of GWAS and eQTLs was significantly associated with the bias (p<2E-16, Wald test in linear regression). A smaller expression sample tended to result in a larger bias (Figure S8a and S8b). In addition, we found that the bias was also associated with the causal effects (p<2E-16, Wald test in linear regression), and there were larger biases at larger causal effects. A linear regression model with 2-degree polynomial of the sample-size ratio and estimated causal effect can correct the estimation bias for IVW MR method and ML-based MR method (Figure S8c, S8d and Figure S9). For binary phenotypes, we also observed similar patterns in the deviation of estimated causal effects from the true effects. The absolute values of estimated effects were also smaller than that of the true effects in general (Figure 4c, S10-13).

**Figure 4:**
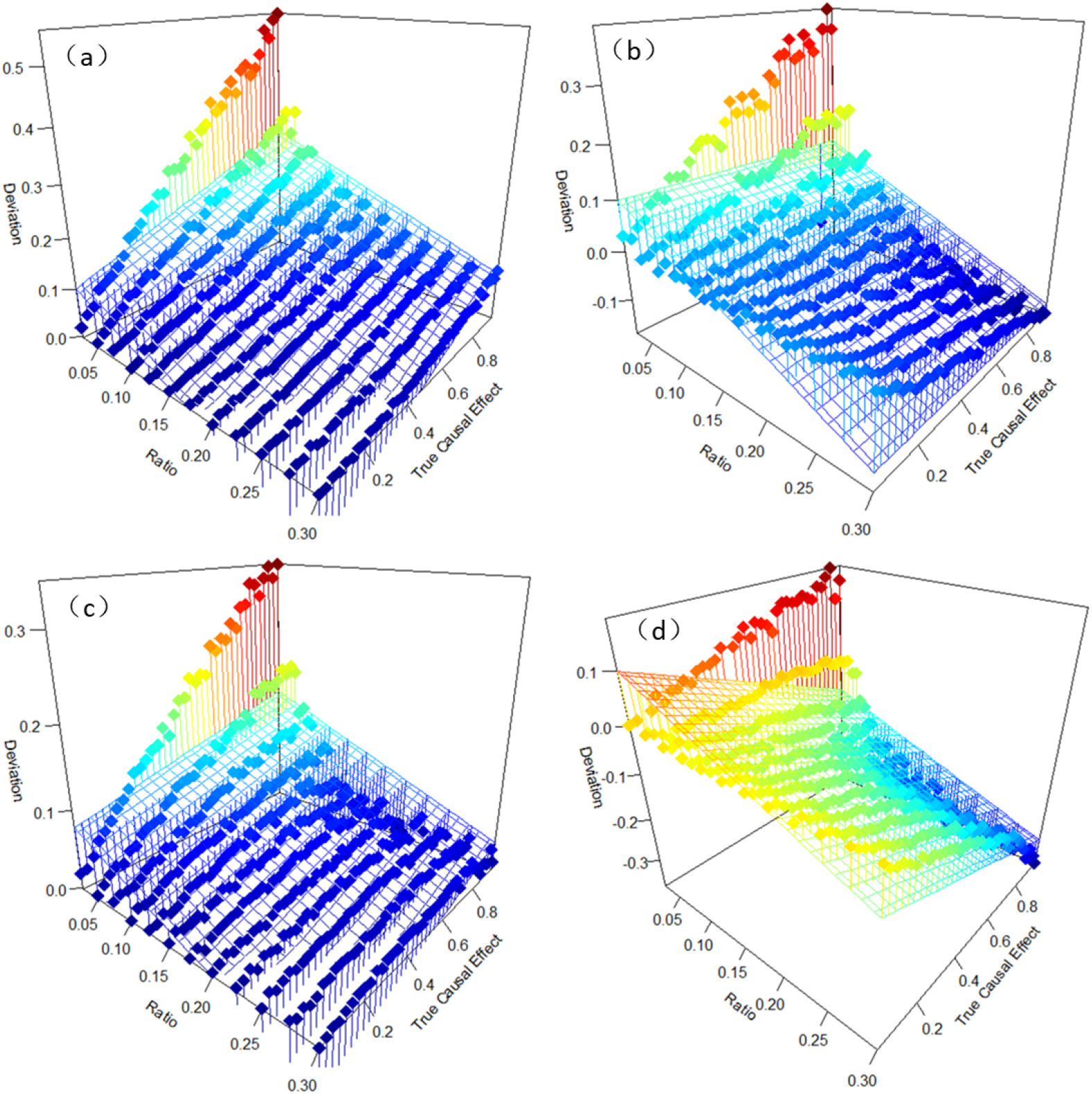
Estimation deviation of causal effect on different types of phenotypes by ML-based MR a) estimation deviation causal effect for continuous phenotype with full set of correlated IVs, b) estimation deviation causal effect for continuous phenotype with LD-pruned IVs, c) estimation deviation causal effect for binary phenotype with full set of correlated IVs, d) estimation deviation causal effect for binary phenotype with LD-pruned IVs. The ratio refers to the size ratio of eQTLs expression sample and GWAS sample. The deviation refers to the difference between estimated effects and true effects.

Two valid MR methods (i.e. ML-based MR and median-based MR) with moderately LD-pruned IVs also showed biased estimation. For continuous phenotypes, the absolute values of estimation were also smaller than the true effects (Figure 4b and S14), which was similar to that based on full set of IVs. The smaller sample size ratio of expression sample to GWAS sample led to larger estimation deviation (p<2E-16, Wald test). In addition, the deviations also increased generally as true effects became larger (*p*<2E-16, Wald test). Among the two methods, the ML-based MR method showed lower deviation than median-based MR method. For instance, when the expression sample size was relative large (say, n>1500) and true effect was moderate (say, ±0.2), the estimate causal effect was very close to the true effects, ∼±0.18 (Figure 5a and 5b). However, for binary phenotypes, the absolute values of most estimations were larger than the true effects when the sample size ratio was not small, say >10%, corresponding to 2000 subjects (Figure S15, 5c and 5d). However, in a scenario close to practical scenario in which the sample size of eQTLs was 500 (corresponding to the ratio 2.5:100), the estimated effect sizes by ML-based MR method and median-based MR method were 20% and 40% smaller than of the true effects respectively.

**Figure 5:**
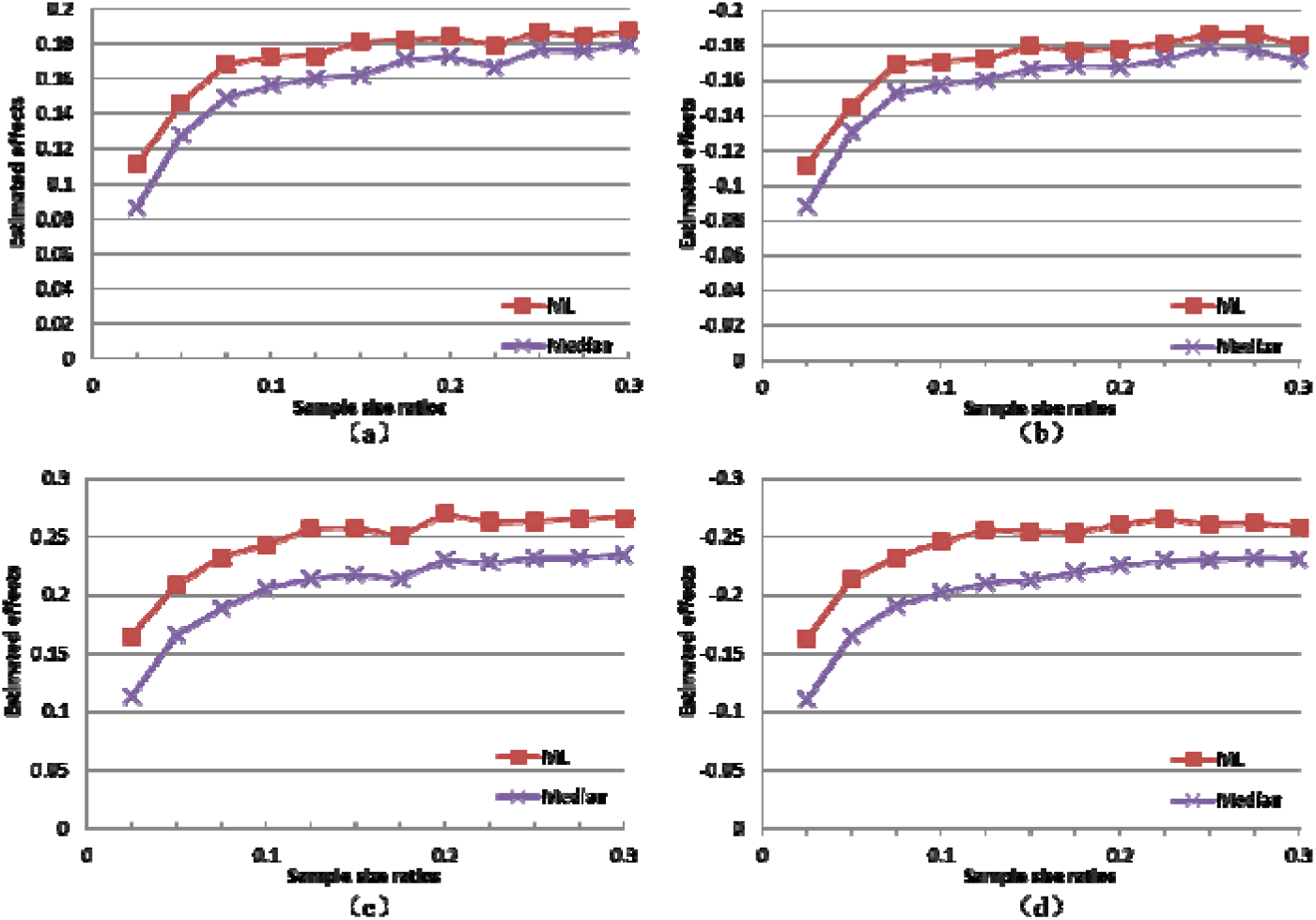
The estimated causal effects by different methods. a) the true effect was 0.2 for continuous phenotypes, b) the true effect was -0.2 for continuous phenotypes, c) the true effect was 0.2 for binary phenotypes d) the true effect was -0.2 for binary phenotypes. Each data point was an averaged estimation in 100 simulated datasets. The outcome phenotype was a continuous phenotype.

### Case studies

#### Schizophrenia

Using GWAS summary statistics of schizophrenia, we investigated how the LD, gene boundary extension and expression type influenced causal gene estimation. Unexpectedly, we found the MR methods using LD matrix with a full set IVs detected much more significant genes than the usage of moderately LD-pruned IVs although they had valid type 1 errors in our simulated data. For instance, the ML-based MR with the full IVs and moderately LD-pruned IVs (threshold *r*=0.5) based on the 1 MB extension and gene level expression detected 663 and 89 significant genes (p<2.5E-6) respectively. Only 34 out of the 663 and 89 significant genes were overlapped. We found 232 of the 663 significant genes’ p-value equal to zero because of unreasonably large estimated causal effects (>1) and/or very small standard error (<1E-3), which may be caused by un-matched LD matrixed. In contrast, the most significant gene among the 89 gene was C4A (causal effect beta= 0.156±0.0145, p=4.45E-27), an well-established causal gene of schizophrenia in recent years^26 27^. It was also the most significant genes prioritized by the median-based MR method (causal effect beta= 0.14±0.008, p=1.9E-60). Therefore, to produce more reliable results, we used the moderately LD-pruned IVs (threshold *r*=0.5) by the ML-based MR and median-based MR (the two methods with valid type 1 error with LD-pruned IVs according to our preceding simulation studies) in the subsequent analyses with the real data.

There were three additional interesting patterns in the MR analysis with schizophrenia GWAS summary statistics. First, the usage of eQTLs based on gene-level expression and transcript-level expression output very different significant gene lists. For example, among the 76 and 65 significant genes respectively (p<2.5E-6) prioritized by the median-based MR method using eQTLs according to transcript-level and gene-level expression with 1 MB extension, only 35 genes (∼50%) were overlapped. Therefore, although the transcript-level expression led to slightly more significant genes, it was still necessary to also use gene-level expression for causal gene estimation. Second, when the gene boundary extension was over 1MB, further extension can moderately increase the number of significant genes. Based on transcript level expression, we found the 2MB upstream and downstream extension only output 1 and 2 more significant genes than 1 MB extension by the ML-based MR method and median-based MR method respectively (Figure 6). When the extension decreased to 100 KB from 1MB, the number of significant genes reduced substantially. For instance, the number of significant genes prioritized by median-based MR method according to transcript-level expression with the two different extension decreased by 44, from 76 to 32. Third, the ML-based MR method detected more significant genes than the median-based MR method. As shown in Figure 4, the ML-based MR method always detected 20 or more significant genes than the median-based MR method at various boundary extensions. The percentage of overlapped significant genes by the two methods were moderate, ∼60% on average. One reason for the difference is that median-based MR method cannot work for genes with less than 3 eQTLs while ML-based MR can do. We also noticed that the p-values of top three genes prioritized by median-based MR method were more significant than that of ML-based MR method, which was consistent with the power evaluation results in the preceding simulation.

**Figure 6:**
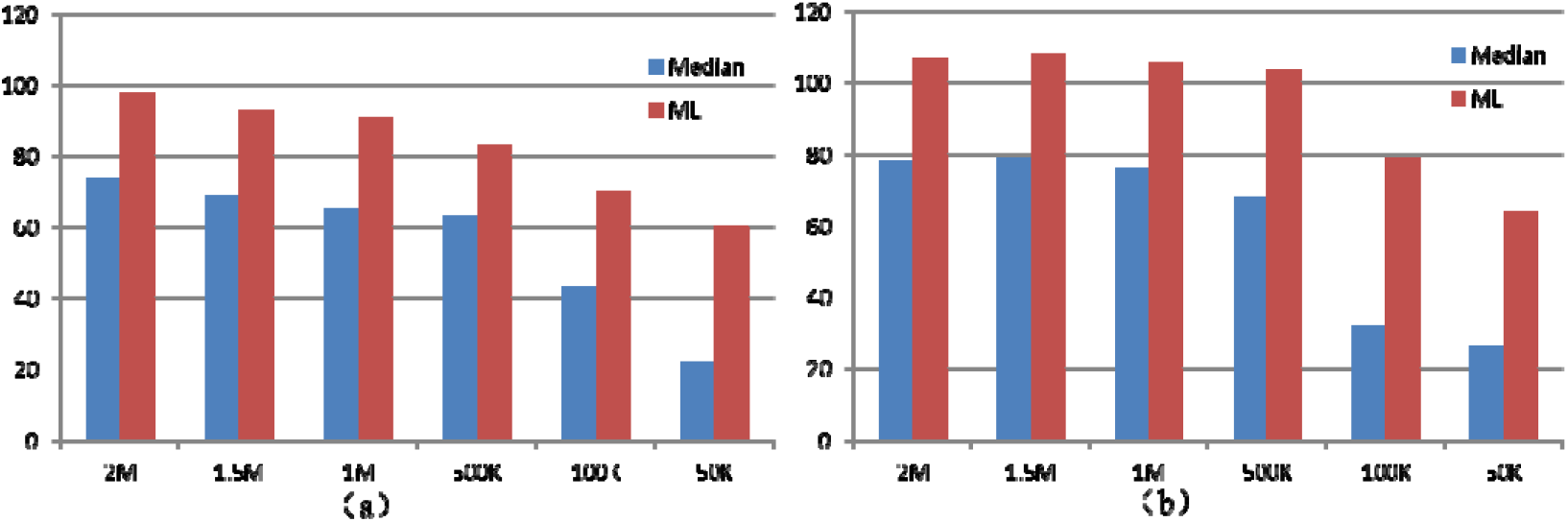
Number of significant causal genes for schizophrenia. a) MR analyses based on gene-level expression; b) MR analyses based on transcript-level expression. The X axis denotes the length of gene boundary extension to cover more potential regulatory eQTLs.

We then looked into details of significant genes prioritized by the two valid MRs based on the 1MB extension and LD-pruned IVs (*r*=0.5). The ML-based MR method and the median-based MR method based on gene-level expression detected 89 and 64 significant genes (p<2.5E-6) respectively. Among these significant genes, 25 and 20 genes had at least one hit paper that mentioned the gene symbol and schizophrenia in abstracts in PubMed (See details in Excel Table S1). Eleven genes, including 4 MHC genes (C4A, HLA-B, HLA-C, and HLA-DQA1) and 7 non-MHC genes (NT5C2, FURIN, GLT8D1, NOTCH4, GATAD2A, NEK4, and MAPK3), had over 5 hit papers. The MHC gene C4A was prioritized as the top gene by both MR methods according to the p-values, which is also a well-known causal gene of schizophrenia ^28^. Figure 7a visualizes relationship of the association signals in a scatter plot. The estimated causal effects 0.156±0.0145 (p=4.45E-27, by ML-based MR) suggested its over expression increased the risk of schizophrenia, which was consistent with a very recent finding ^28^. Among the 8 non-MHC genes, the most significant one was GATAD2A, *p*=9.18E-24 by ML-based MR method. GATAD2A encodes GATA zinc finger domain and is a transcriptional repressor. It has been suggested as an associated gene of schizophrenia ^29^. The estimated negative causal effect (causal effect beta= -0.093±0.009) suggested the gene’s expression may have protective effect on schizophrenia (Figure 7b). Consistently, Huckins et al also estimated a negative causal effect of this gene on schizophrenia ^30^. The second most significant gene by median-based MR method was VARS2 (*p*=3.26E-38). VARS2 encodes a mitochondrial aminoacyl-tRNA synthetase. Fromer et al also prioritized it as an important associated gene of schizophrenia ^31^. Nevertheless, most of the estimated causal genes (Excel Table S2) are new for schizophrenia and are subject to validation in independent samples or by biological experiments.

**Figure 7:**
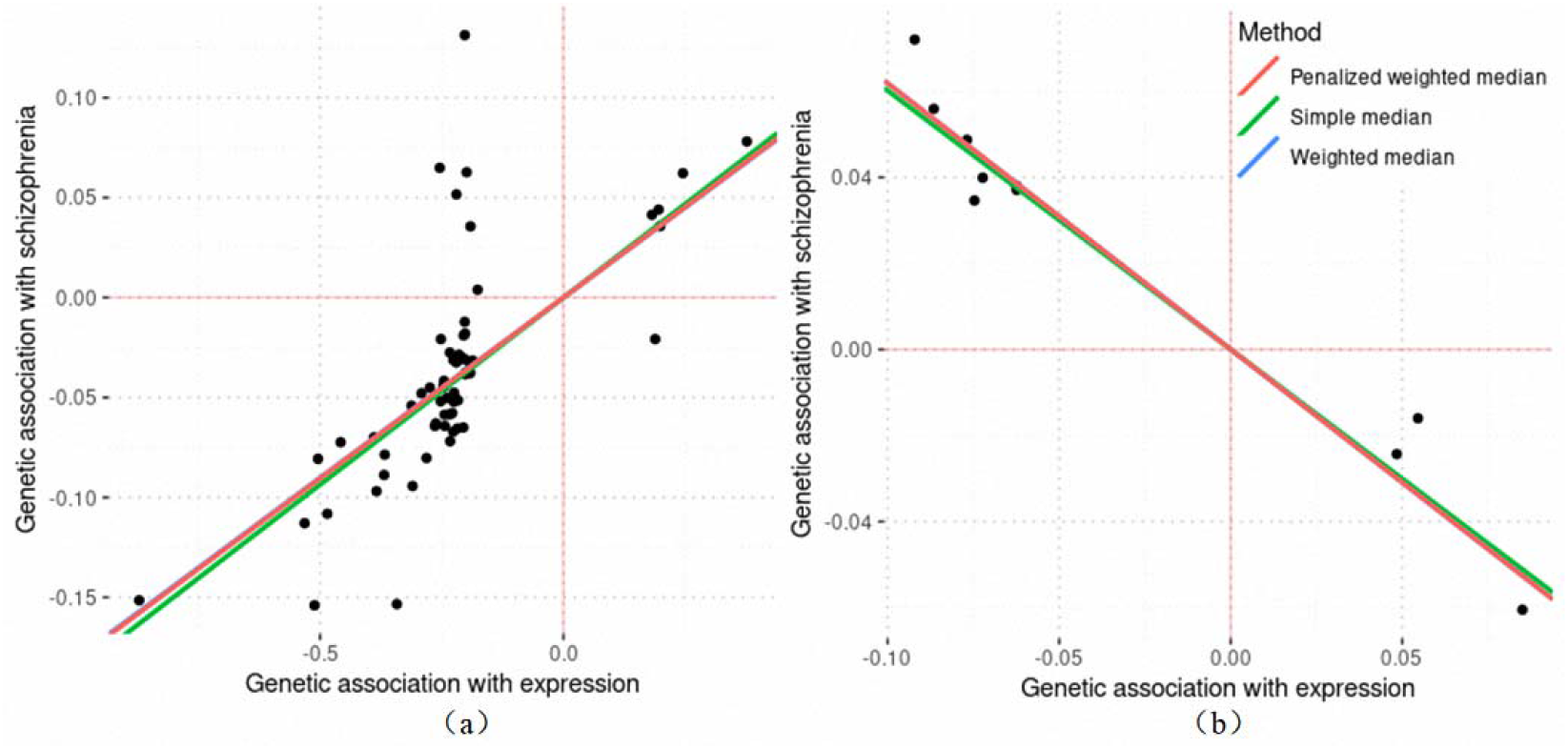
Scatter plots of genetic association with gene expression in schizophrenia. a) eQTL and GWAS summary statistics in C4A; b) eQTL and GWAS summary statistics in GATAD2A. A dot on the plots denotes an IV.

As a comparison, based on transcript-level expression, the ML-based MR method and the median-based MR method detected 106 and 76 significant genes respectively (p<2.5E-6 after Bonferroni correction for the number of transcripts within a gene). Among these significant genes, 36 and 28 genes had at least one hit paper in PubMed database respectively (Excel Table S3). Thirteen genes had over 5 hit papers, in which 4 genes were in the MHC region. Compared to significant genes according to gene-level expression, five out of the 13 genes were unique according to transcript-level expression, namely, MAPT, HSPA1B, SDCCAG8, TSNARE1 and PCNT. The C4A gene remained the most and the fourth most significant gene by the median-based and ML-based MR methods respectively. Among the 13 genes, SDCCAG8 had third most significant p-value by the ML-based MR method. SDCCAG8 encodes a centrosome-associated protein that may be involved in organizing the centrosome during interphase and mitosis. Its significant transcript was ENST00000463042, *p*=5.80E-11, in which the higher expression had a protective effect on the risk of schizophrenia (causal effect beta= 0.135±0.021). Consistently, William and Murray found predicted expression of SDCCAG8 was associated with higher risk of schizophrenia in the DLPFC ^32^. Similar to the estimation based on the gene-level expression, many genes with fewer and even no hit papers might be also promising causal genes (Excel Table S4). For example, GNL3 only had two hit papers. It was the second and fourth most significant gene by the ML-based MR method and median-based MR method respectively. Its isoform ENST00000394799 achieved a p-value *p*= 1.65E-39 (causal effect beta= 0.115±0.009) by ML-based MR method. A very recent study suggested that eQTLs may regulate expression of GNL3 and further influence risk of psychiatric disorders ^33^.

#### Bipolar disorder

The general patterns of significant genes in bipolar disorder were similar to that of schizophrenia in general. The 1MB gene boundary extension seemed also sufficient to cover most cis-eQTL to detect most significant casual genes. The number of significant genes based on 1MB extension was very similar to that of 2MB extension, e.g., 55 vs. 57 by the ML-based MR method with gene-level expression. A further decrease in the bounder to 100KB substantially reduced the number of significant genes (Figure 8). The ML-based MR method still detected more significant genes than the median-based MR method. However, the difference in the number of significant genes between the two methods in this disease was much larger than that of schizophrenia, e.g., 55 vs. 19 by the ML-based MR method and median-based MR method respectively with 1 MB extension based on gene-level expression. The transcript-level expression output smaller number of significant genes than the gene-level expression in bipolar disorder. Again, the percentage of overlapped significant genes based on the two different levels of expression were not high either, <50%. Finally, compared to the above schizophrenia results, the number of significant genes in the bipolar disorder dataset was smaller. For example, given 1MB extension and gene level expression, the ML-based MR method detected 55 significant genes for bipolar disorder while it detected 91 significant genes for schizophrenia. Probably, this was because the GWAS sample of bipolar disorder was smaller than that of schizophrenia, 46,582 vs. 117,498.

**Figure 8:**
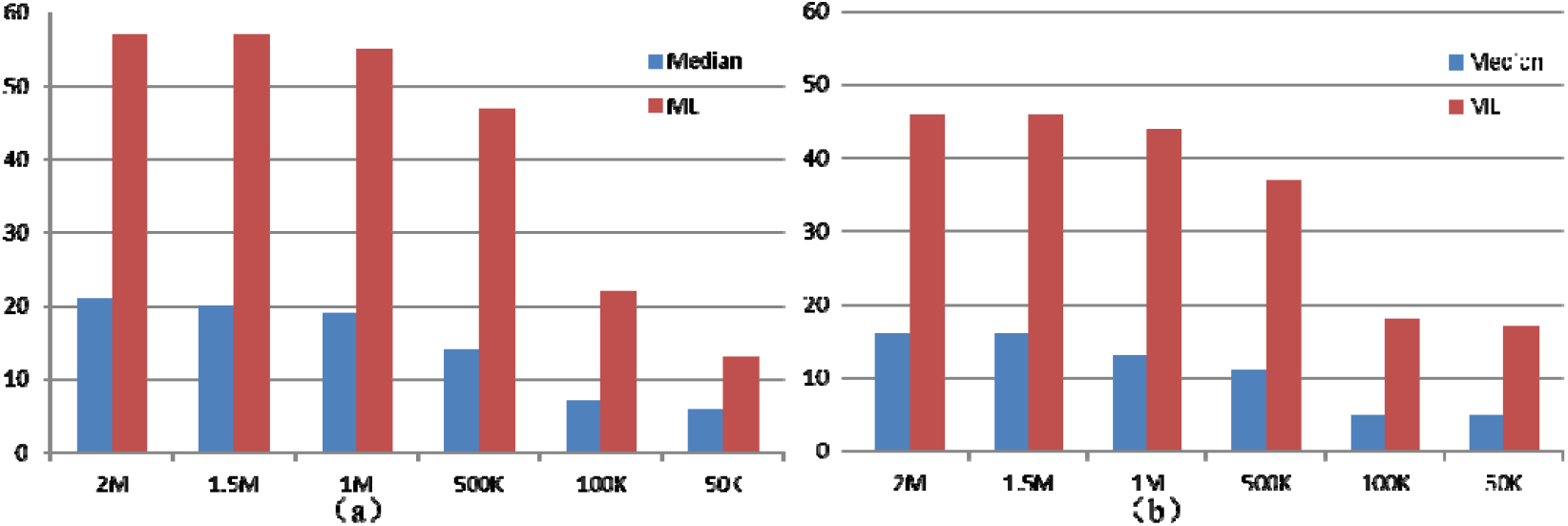
Number of significant causal genes for bipolar disorder. a) MR analyses based on gene-level expression; b) MR analyses based on transcript-level expression.

The ML-based MR method and the median-based MR method using gene-level expression and 1MB gene boundary extension detected 55 and 19 significant genes respectively. Probably because bipolar disorder is less studied than schizophrenia, among the significant genes, the genes with one or more hit papers in PubMed database (9 and 4 for the two methods) were much fewer than that of schizophrenia (Table S5). Only three genes (C4A, FADS1 and NEK4) among all the significant genes had 3 or more hit papers. Similar to schizophrenia, C4A was also the most significant causal gene of bipolar disorder, *p*=3.24E-15, by ML-based MR. But its estimated effect in bipolar disorder was smaller than that of schizophrenia, 0.048±0.006 vs. -0.16±0.015 respectively. Melbourne et al found C4A mRNA expression in peripheral blood mononuclear cells could predict the presence and severity of delusions in bipolar disorder with psychosis ^27^. FADS1 was the third most significant genes according to the ML-based MR, *p*=8.74E-15. It encodes a member of the fatty acid desaturase gene family. Ikeda et al detected a significant locus close to this gene associated with bipolar disorder in a Japanese sample ^34^. They also found the top association SNP (rs28456) was a strong eQTL with causal effect on FADS1 in brain regions multiple from independent datasets by SMR method. The association between this gene and bipolar disorder was also found in a Chinese sample ^35^. The ML-based MR estimated its over expression had protected effect on bipolar disorder, causal effect beta= -0.14±0.018. NEK4 encodes a serine/threonine protein kinase for normal entry into replicative senescence. Its overexpression is estimated to increase risk of bipolar disorder, causal effect beta=0.186±0.028 by ML-based MR method. Recently, Yang et al showed biological evidences for the causal effects of NEK4 and adjacent genes on psychiatric disorders ^33^. Nevertheless, most of the genes (Excel Table S6) are new for bipolar disorder and are subject to further validation.

The ML-based MR method and the median-based MR method based on transcript-level expression and 1MB gene boundary extension detected 44 and 13 significant genes respectively. Only 6 and 3 of the significant genes had one or more hit papers in PubMed database respectively (Table S7). There were four genes with 3 or more hit papers, FADS1, C4A, GNL3 and S100B. The latter 2 genes were overlooked by the analysis based on gene-level expression. C4A were the fourth and the second most significant causal gene of bipolar disorder by ML-based MR method and the median-based MR method. GNL3 was the fourth most significant causal gene by ML-based MR method. It encodes the G protein nucleolar 3. Interestingly, two transcripts of this gene were estimated to have opposite casual effects, ENST00000394799:0.16±0.02 and ENST00000468146:-0.26±0.048. This may be the reason why the MR analysis based on gene level expression (which was the averaged expression of transcripts) overlooked this gene. A very recent study showed that GNL3 knockdown and overexpression led to aberrant neuronal proliferation and differentiation ^36^. S100B encodes a member of the S100 family of proteins containing 2 EF-hand calcium-binding motifs. Serum S100B protein has been used as an important biomarker of bipolar disorder ^37^. Serum S100B protein’s expression level was increased in patients with bipolar disorder ^38^ while decreased after treatment in bipolar patients in a manic phase ^39^. Consistently, its most significant transcript ENST00000397648 had a positive effect on bipolar disorder, causal effect beta=0.20± 0.035, *p*=4.84E-9. Our analysis further suggested that S100B might be more than a biomarker of this disease. Many genes (See details in Excel Table S8) without hit paper were also promising causal genes for bipolar disorder. For example, GOLGA2P7 was the most significant gene by the median-based MR method with a causal effect beta=0.4±0.06 at its transcript ENST00000559668, *p*=3.43E-11. GOLGA2P7 is a pseudogene of GOLGA2. A very recent study reported the association between GOLGA2P7 and bipolar disorder ^40^. The significant causal effects of other genes and transcripts are subject to be validated.

## Discussion

This is the first study so far, to our knowledge, evaluating the performance of six widely-used MR methods original designed for epidemiological studies in a new type of application — inferring causal genes with a small-scale expression data and a large-scale of GWAS data in the post-GWAS era. Through systematical evaluation study with both simulated and real data, we prioritized ML-based MR out of the six methods as the most robust and suitable approach. When the IVs are correlated, the ML-based MR had valid type 1 error and sufficient power in simulated data. The ML-based MR method also worked with LD-pruned IVs according to a small or moderate cutoff *r*<0.5. More importantly, it was able to generate a reasonable number of significant genes and “rediscover” many known susceptibility genes of schizophrenia and bipolar disorder in real datasets. The median-based MR method could also work with LD-pruned IVs but produce fewer significant genes in real data analysis. The mode-based MR method and contamination mixture MR method cannot model LD between IVs and had inflated type 1 errors while IVW MR method had inflated type 1 error with moderately LD-pruned IVs. The MR-Egger method had low power with LD-corrected IVs and had inflated type 1 error with moderately LD-pruned IVs. Moreover, we also explored a best practice pipeline for causal genes inference with GWAS summary statistics and eQTLs. We found the 1MB gene boundary extension may be sufficient to cover most regulatory variants for a powerful inference. Probably due to different efficiency in detecting eQTLs, it is necessary to use eQTLs of both gene- and transcript-level expression for the inference. In real data analysis, as the MR methods are sensitive to noises in LD matrix, the usage of LD-pruned IVs will lead to more reasonable estimation than the full set IVs. Based on the prioritized methods (the ML-based MR and median-based MR) and the optimized pipeline, a powerful tool integrated with whole-genome eQTLs at over 50 tissues or cell types, named MACG, was developed for causal genes inference by using GWAS summary statistics, available at http://pmglab.top/kggsee.

This is also the first study so far to investigate how the expression of different levels (gene vs. transcript) influenced the causal genes estimation. We found only less than 50% of significant genes based on the two different levels of expression were overlapped. Theoretically, there are pros and cons for the usage of different levels of expression. As the amount of gene expression is larger than that of the transcript, the former will lead to more significant eQTLs for the inference. Consequently, the MR methods would be more powerful to detect some causal genes with homogeneous expression. However, the gene-level expression essentially is just an averaged expression of multiple transcripts. When there is large heterogeneity among the transcripts’ expression, the averaged expression will neutralize opposite effects of transcripts (if available) and then decrease the power of MR analysis (see an example at GNL3 for bipolar disorder). With the transcript-level expression, the MR methods will have higher power to detect some risk transcripts of heterogeneous expression. In the real data analysis, the MR methods with transcript level expression rediscovered a known schizophrenia susceptibility gene, SDCCAG8, which was overlooked by MR analysis based on gene level expression. SDCCAG8 had 2 transcripts and only one of transcripts achieved a significant p-value 5.8E-11. Therefore, we recommend using both levels of expression to generate eQTLs, if available, for the causal genes inference.

Our study is different from existing methods integrating gene expression for genetic mapping. First, our framework, MACG, is different from an alternative MR approach, SMR ^11^. It only used a single IV (usually the top SNP) in the analysis, although for the same purpose. In contrast, MACG uses multiple variants simultaneously to infer causal genes based on MR. We showed that more IVs increased the power to detect the causal genes (Table 1). Barbeira et al. also pointed out that the 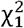 statistics of SMR tended to be deflated under null hypothesis^13^. Second, our framework is also different from some recent multi-variants based integrative association methods, e.g. S-TWAS ^41^ and S-PrediXcan ^13^. These methods were designed to examine association between phenotypes and imputed gene expression in which multiple cis-eQTLs were used to predict gene expression. However, these methods only detected association *per se* but not causation.

Our simulation study suggested that the correlation of IVs had worse influence on performance of the MR methods than the pleiotropy. When full set IVs in LD were used for the MR analysis, the methods that were unable to model the correlation had inflated type 1 error. We found the ML-based MR method and median-based MR method were less sensitive to LD, and thus they had valid type 1 error with moderately LD-pruned IVs approximately. There are at least two reasons why the pleiotropy in IVs had weak effect on the performance of the MR methods. First, for complex phenotypes of polygenic model ^42^, the heritability of individual SNPs for gene expression is unlikely large (<1%) while the sample sizes for eQTLs calculation are usually small or moderate, *n*<1000. Therefore, many pleiotropic SNPs may not survive the hypothesis test on eQTLs. Second, some methods were designed to tolerate moderate pleiotropy. For instance, the median-based MR’s weighted median can provide a consistent estimate if at least 50% of the weight comes from valid IVs ^21^. The random-effects models of ML-based MR may be robust to the pleiotropic IVs to some extent. We found ML-based MR became inflated with pleiotropic IVs when it was carried out under a fixed effect model (Figure S16).

In the proof-of-principle examples, the ML-based MR method and median-based MR method estimated many potential causal genes for schizophrenia and bipolar disorder. Among the significant genes, quite a few genes are known susceptibility genes of the corresponding diseases, e.g., C4A, GATAD2A, MAPK3 for schizophrenia and FADS1, S100B for bipolar disorder. Note that there are also many genes which are new for the diseases. Functional validation of these genes is indispensable to confirm their causality although this is beyond the scope of the present paper. However, we also noticed that some known susceptibly genes of the diseases were not significant in our analysis, e.g., NRG1 for schizophrenia ^43^. This may be related to the power of the MR methods. In our analysis, the expression dataset only had 592 subjects. To reduce weak IVs bias ^23^, we excluded many eQTLs with *p*> 1E-4. For median-based MR method, genes having less than 3 IVs were excluded, which may also overlook some causal genes. In addition, gene expressions are variable in different brain regions and development stages. Our previous study showed that multiple brain regions were highly related to schizophrenia and bipolar disorder besides the frontal cortex ^7^. Due to limited sources of expression data, we only used the dorso-lateral prefrontal cortex regions. Probably, by only using the gene expression in dorso-lateral prefrontal cortex, other related brain regions’ causal genes may be overlooked by this study..

It is unexpected that the consideration of LD correlation in real data analysis led to numerous false positive discoveries although it worked well in simulated data. We compared the details in estimation with LD correction with that of LD pruning. A main problem was that the former usually had extremely small SE of the estimated causal effect than that of the latter, e.g., <1E-4 for former vs. >0.01 for latter. The calculation of SE with LD correction includes inversion and multiplication of the LD matrix. In this study, we used the 1KG genotypes as a proxy dataset to approximate LD matrix of the large-scale GWAS meta-analysis for schizophrenia and bipolar disorder. It is very likely that there are noises from the LD matrix due to smaller sample sizes and incompletely matched ancestry. Therefore, the usage of noised LD matrix may lead to a very small SE, which subsequently resulted in an inflated *z* score or type 1 error. In the simulation, we used the sample itself rather than a proxy to calculate the LD matrix so that the estimated SEs were correct. In practice, it is infeasible to merge all genotypes of a large GWAS meta-analysis study to calculate the LD matrix as we did in simulation. Therefore, ML-based MR method and median-based MR method had valid type 1 error even with moderately LD-pruned IVs according to reference sample, and thus may be more realistic options for inferring causal genes.

A limitation of the present study is lack of validation for new causal genes. We did not validate the estimated significant causal genes of schizophrenia and bipolar disorders in additional samples or biological experiments. Therefore, it is unknown how many significant genes are true causal genes although we rediscovered a number of known susceptibility genes for schizophrenia and bipolar disorders respectively. Most of the significant genes are recommended to be investigated and validated by independent genetic samples or functional experiments. Therefore, we shared the significant genes and their causal effects in the supplementary Tables (S1-4), which may be valuable reference of candidate causal genes for follow-up studies in the future.

## Methods and Materials

### Mendelian randomization methods

The following six popular MR methods were investigated by comparing their type 1 error rates and powers for causal gene inference. The first three MR methods can be used to adjust for correlation of IVs while the other three MR methods cannot do. In addition, the last four methods were designed to account for horizontal pleiotropy in IVs by different strategies. The R codes of the six methods implemented in the MendelianRandomization R package (version 0.4.2)^44^ were called by MACG for analyses in the present study. The following is a brief description of the six MR methods.

#### The inverse-variance weighted (IVW) MR

The causal effect was estimated by using generalized weighted linear regression with the associations with the phenotype and with the gene expression, in which the intercept was set to zero and weights were the inverse-variances of the associations with the phenotype ^20^. The genotypic correlations of variants (known as IVs) were used to correct relatedness of associations with phenotypes. The random-effects model with correlated IVs was adopted, in which the main R code was “mr_ivw(mr_input, correl=TRUE, model=‘random’)”. In other testing scenarios, the LD pruned IVs according to various LD cutoffs were input into a random-effects model, in which the R code was “mr_ivw(mr_input, penalized=TRUE, robust=TRUE, weights=‘delta’, model=‘random’)”.

#### The maximum-likelihood (ML) based MR

The causal effect was estimated by a likelihood function of the bivariate normal distribution for the associations of each genetic variant with gene expression and with the phenotype. The mean of the association with the phenotype was taken as the mean association with the expression multiplied by the causal effect parameter. A random-effects model was also adopted. Compared to the IVW MR method, this MR method may have advantages of incorporating uncertainty in the genetic associations with the expression. The main R code was “mr_maxlik(mr_input, correl=TRUE, model=‘random’)”. LD pruned IVs were tested and the used main R code was “mr_maxlik(mr_input, model=‘random’)”.

#### MR-Egger method

MR-Egger method was proposed by Bowden et al (2015) ^25^ for the scenario where genetic variants were not all valid IVs. MR-Egger intercept test can be used to estimate directional pleiotropy. This method is supposed to provide an unbiased test for a causal effect when the instrument strength independent of direct effect (InSIDE) assumption holds ^25^. The main code of this method in the MendelianRandomization R package was “mr_egger(mr_input, correl=TRUE)”. LD pruned IVs were tested by the R code “mr_egger (mr_input, penalized=TRUE, robust=TRUE)”.

#### Mode-based MR

The mode-based estimation (MBE) MR method was proposed by Hartwig et al ^22^. It takes the variant-specific ratio estimates from each genetic variant in turn, and calculates the model by a kernel-smoothed density out of the ratio estimates. Like the MR-Egger, the method was developed to relax the IV assumptions. As it cannot be used to correct the relatedness between IVs, we tested the method with the full set of IVs and LD pruned IVs separately with the same R code, “mbe(mr_input, stderror = ‘delta’)”.

#### Median-based MR

The weighted median MR method calculated the median of the ratio IV and used weighted median estimator for combining data on multiple genetic variants into a single causal estimate ^21^. While working for moderately invalid IVs, it also has greater robustness to individual genetic variants with strongly outlying causal estimates compared other MR methods. It also cannot be used to correct the relatedness between IVs. The full set of IVs and LD pruned IVs were tested separately with the same R code “mr_median(mr_input)”.

#### Contamination mixture MR

The contamination mixture MR method was proposed by Burgess et al (2020) ^18^. It was built on a likelihood function of a product of two component mixture normal distributions for valid instruments and invalid instruments respectively. The contribution to the likelihood for each genetic variant as a valid instrument and as an invalid instrument was calculated respectively. The obtained contribution was then compared to configure valid and invalid instruments that maximize the likelihood for the given value of the causal effect. The detailed model and algorithm can be seen in Burgess et al (2020) ^18^. As it cannot be used to correct the relatedness between IVs, the full set and LD pruned IVs were tested separately by the same main R code “mr_conmix(mr_input, CIMin = -1, CIMax = 5, CIStep = 0.01)”.

### Summary data–based Mendelian randomization (SMR)

We also compared the above multiple eQTLs based MR methods with a single eQTL based MR method, SMR^11^. SMR was designed to infer causal genes of a phenotype also by using GWAS summary-level data and eQTLs. The causal effect is estimated by a ratio of coefficients in the least squares regressions and the sampling variance of the estimate is approximated by the Delta method. A χ^2^ statistics is then built to test the causality of gene expression to a phenotype. In our analysis, the summary statistics of eQTLs and phenotype associated variants calculated from simulated data were formatted and input into the tool SMR (version 1.03) with default parameters for the causality test by the χ^2^ statistics. The test p-values were extracted from the output results for the power comparison.

### Simulation of genotypes and phenotypes

To investigate the statistical type 1 error, power, and accuracy of the MR methods for inferring causal genes, extensive computer simulations were performed. A 3MB region which mimicked the length of highly expanded gene was randomly drawn from human genome. In the EUR panel of 1000 Genomes Project ^45^, this region contains 2371 common variants (MAF>0.01). Genotypes of the variants were simulated given allelic frequencies and LD correlation matrix in the according to the HapSim algorithm ^46^. The genotypes were encoded by the number of alternative alleles, *s*. The SNPs’ genotypes (*s*) contributing to the phenotypes were then standardized as, 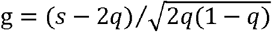, where *q* was the allele frequency of alterative allele. Phenotypes were simulated under a polygenic model of random effect ^42^. One hundred twenty-one independent variants were extracted from the 2371 variants according to stringent LD pruning (r^2^<0.05). Given the heritability (*h*^2^) of a trait and the number of independent causal variants (*m*), the effect size of a variant followed a normal distribution N(0, *h*^2^/*m*). For gene expression, it was assumed m_1_ independent causal variants (total heritability 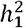) and m_c_ independent pleiotropic variants (total heritability 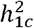) had effect size β_1,*i*_ and *α*_1,*i*_ on gene expression respectively. There were no overlapped variants between the m_1_ and m_c_ variants. In addition, common confounding factors also contributed to *γ*_1_ fraction to the gene expression. Therefore, the expression of a gene was simulated according to the formula:

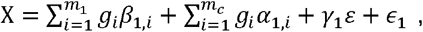

where 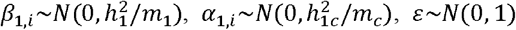 and 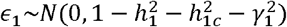.

For a phenotype, it was assumed m_2_ causal variants (total heritability 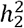) and m_c_ pleiotropic variants (total heritability 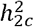) had effect size β_2,*i*_ and *α*_2,*i*_ on the phenotype respectively. In addition, the same common confounding factors also contributed to *γ*_2_ fraction to the phenotype. The above gene expression also contributed δ to the phenotype. The phenotype value was simulated according to the formula:

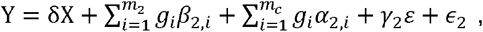

where 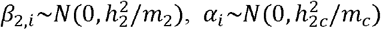 and 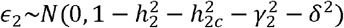.

Here Y was a continuous phenotype. For a binary phenotype, a cutoff *t* was set according to a given disease prevalence *K* under standard normal distribution and the liability threshold model ^47^. Subjects with simulated Y values ≥*t* were set as patients and others were set as normal controls. A large population of 50 million subjects as simulated for each parameter setting. Subjects (N=20,00) were randomly drawn (sampling without replacement) from the large population to form samples for association analysis by linear regression at the gene expression and phenotype respectively. Similar to real situations in practice, there were no overlapped subjects in the samples for the two association analyses. Summary statistics of the association analyses were input to MACG for the MR analysis to infer causal genes.

Fo r binary phenotype, both logistic regression and contingency table analysis were used to calculate the odd ratio (OR) and SEs. The logarithm of OR was converted into effect size under a liability threshold model ^47^. Under the liability threshold model for a binary phenotype, the OR of variants was converted to the equivalent effect size estimation on the liability level. According to Gillett et al. ^47^, the liability level effect size can be approximated by 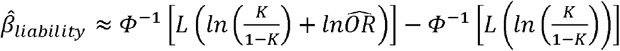 and the variance is 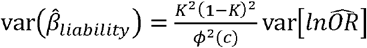, where *ϕ* and *ϕ* are c.d.f and p.d.f of the standard normal distribution respectively, *K* is the disease prevalence, *L* is the standardized logistic function and *c* is the corresponding quantile for the cumulative probability of K under the standard normal distribution.

### Validation on Real datasets

#### GWAS summary statistics dataset

We download real GWAS summary statistics data of two common psychiatric disorders (schizophrenia and bipolar disorder) from public database for further validation. The summary statistics of schizophrenia were derived from a large GWAS meta-analysis [Ref], in which there were 53,386 European cases and 77,258 European controls. The summary statistics of bipolar disorder were also derived from another large GWAS meta-analysis ^48^, in which there were 20,352 European cases and 31,358 European controls. The combined ORs and SEs from meta-analysis were used for the validation analysis. The details of sample sources and data quality controls methods can be seen in the original papers ^49 48^. Genotypes in CEU panel from 1000 Genomes Project ^45^ were used to correct for relatedness of the summary statistics.

### eQTL summary statistics dataset

The summary statistics of eQTLs were calculated based on genotypes and dorsolateral prefrontal cortex gene expression downloaded from CommonMind Consortium. We did not use GTEx eQTLs because there were less than 200 samples for most brain regions (V8). To increase overlap of eQTLs with GWAS summary statistics, the imputed genotypes were used after quality control with INFO score >0.5. After matching genotype data and expression data (in files cmc_isoform-adjustedsva-datanormalization-includeancestry-adjustedlogcpm.tsv.gz and cmc_gene-adjustedsva-datanormalization-includeancestry-adjustedlogcpm.tsv.gz), 527 subjects were retained. To reduce effects of confounding factors, a linear regression model was built in which co-variables (such as disease status, age, sex and so on) were used to adjust the expression. The adjusted gene expression values were used to estimate the effect size and SEs of an eQTL in a simple linear regression model. In the model, the alleles were encoded as 0, 1, and 2, which were in accordance with the coding for the GWAS summary statistics. The allele frequencies were used to assist to flip discordant alleles of variants between the eQTL and GWAS datasets.

## Description of Supplemental Data

Figures S1–S17, Tables S1–S8

## Acknowledgements

This work was funded by National Natural Science Foundation of China (31771401 and 31970650), National Key R&D Program of China (2018YFC0910500), Science and Technology Program of Guangzhou (201803010116).

